# Plant age-dependent dynamics of annatto pigment (bixin) biosynthesis in *Bixa orellana* L.

**DOI:** 10.1101/2022.12.28.522146

**Authors:** Kleiton Lima de Godoy Machado, Daniele Vidal Faria, Marcos Bruno Silva Duarte, Lázara Aline Simões Silva, Tadeu dos Reis de Oliveira, Thais Castilho de Arruda Falcão, Diego Silva Batista, Marcio Gilberto Cardoso Costa, Claudete Santa-Catarina, Vanildo Silveira, Elisson Romanel, Wagner Campos Otoni, Fabio Tebaldi Silveira Nogueira

## Abstract

Age affects the production of secondary metabolites, but how developmental cues regulate secondary metabolism remains poorly understood. Annatto *(Bixa orellana* L.) is a source of bixin, an apocarotenoid used in the world’s food industry worldwide. Understanding how age-dependent mechanisms control bixin biosynthesis is of great interest for plant biology and for pharmaceutical, cosmetic, and textile industries. Here, we used genetic and molecular tools to unravel the role of the annatto age regulated miRNA156 (miR156) targeted *SQUAMOSA PROMOTER BINDING PROTEIN LIKE (BoSPL)* genes in secondary metabolism. Low expression of several *BoSPL* genes in miR156 overexpressing annatto plants (OE::156) impacted leaf ontogeny, reducing bixin production and increasing abscisic acid (ABA) levels. Modulation of *BoCCD4;4* and *BoCCD1* expression, key genes in lycopene cleavage, was associated with diverting the carbon flux from bixin to ABA, whereas upregulation of *lycopene β cyclase* genes implies the xanthophyll biosynthetic pathway acted as a carbon sink in OE::156 plants. Proteomic analyses revealed low accumulation of most secondary metabolite-related enzymes in OE::156 plants, suggesting that miR156 targeted *BoSPLs* are required to activate several annatto secondary metabolic pathways. Our findings suggest that carbon flux in *B. orellana* OE::156 leaves was redirected from bixin to ABA production, indicating an age-dependent leaf dynamics of bixin biosynthesis. Importantly, our study opened a new venue to future annatto breeding programs aiming to improve bixin output.

## Introduction

The vegetative-phase change pathway controls the transition between juvenile and adult phases in plants, affecting several fundamental metabolisms, including the production of secondary metabolites (Li *et al*., 2020). It is well known that secondary metabolites vary at different stages of the plant life cycle, but how endogenous developmental cues regulate secondary metabolism remains poorly understood. Leaf age and developmental stage affect the production of secondary metabolites (Vazquez-Leon *et al*., 2017; Li *et al*., 2016). For instance, the biosynthesis of some monoterpenes and sesquiterpenoids begins as early as in the first cotyledon in *Melaleuca alternifolia* (Russell and Southwell, 2002). Synthesis and accumulation of essential oil of *Cinnamomum verum* initiates in young leaves (Li *et al*., 2016). Nevertheless, the synthesis of compounds associated with the sabinene hydrate–terpinen-4-ol–γ-terpinene pathways are produced at later stages of leaf development (Russell and Southwell, 2002). In terms of defense, total phenolics, alkaloids and cyanogenic glycosides increase in some species when plant ages (Elger *et al*., 2009). Localized on the surface or inside the leaves, secretory structures such as nectaries, resin ducts, secretory vesicles, salt glands, oil cells and secretory trichomes, are frequently one of the main sites for synthesis and accumulation of secondary metabolites (Figueiredo *et al*., 2008). Therefore, distinct ontogenetic stages of leaves may affect the concentration of various primary and secondary plant products (Verma and Shukla, 2015). Although leaf ontogenetic changes are genetically programmed, they are not necessarily synchronized with whole-plant ontogenetic changes. New leaves are continuously produced throughout the vegetative phase, and long-lasting plants such as trees may represent a complex mosaic or a gradient of leaf tissues at different ontogenetic phases of development. These ontogenetic differences may impact the production of secondary metabolites in the plant as a whole. However, little is known about the molecular mechanisms underlying the age-dependent control of the secondary metabolism in non-model plants, such as trees.

Popularly known as achiote tree or annatto, *Bixa orellana* L. (Bixaceae) belongs to the order Malvales, whose representatives include cocoa, hibiscus, and cotton (The Angiosperm Phylogeny Group II, 2003; 2009). Annatto is a source of the secondary metabolite bixin, an apocarotenoid commonly used in the world’s food industry worldwide (Ramamoorthy *et al*., 2010; Vilar *et al*., 2014). In young leaves and seeds of *B. orellana*, apocarotenoids as bixin are derived from the oxidative cleavage of carotenoids, through the action of specific enzymes previously identified such as lycopene cleavage dioxygenase (BoLCD), carotenoid dioxygenases (BoCCDs) (Soares *et al*., 2011), aldehyde dehydrogenases (BoALDHs), and SABATH family methyltransferase (BoSABATH) (Cárdenas-Conejo *et al*., 2015). Bixin pathway may share common enzymes and substrates with other carotenoid-derived compounds, such as abscisic acid (ABA) (Cazzoneli and Pogson, 2010). Bixin is non-toxic, as it is obtained from a natural source, and does not alter the nutritional value of foods (Vilar *et al*., 2014). Annatto also contains a hydrophilic natural dye, norbixin (Giuliano *et al*., 2003; Rivera-Madrid *et al*., 2013), and has broad-spectrum antimicrobial properties (Karmakar *et al*., 2018) provided by other secondary metabolites, such as terpenes, sesquiterpenes, and alkaloids (Santos *et al*., 2021). Understanding the age-dependent underlying mechanisms controlling the biosynthesis of bixin and norbixin is not only of great interest for plant biology, but also for the pharmaceutical, cosmetic, and textile industries (Valério *et al*., 2015).

The main network controlling the transition between juvenile and adult phases includes the temporally regulated microRNA miR156 and its transcription factor targets, members of the *SQUAMOSA PROMOTER-BINDING PROTEIN LIKE (SPL)* gene familym(Cardon *et al*.,1999; Wang *et al*., 2009; Wu *et al*., 2009; Yu *et al*., 2010; Morea *et al*., 2016). The highly conserved miR156/SPL module has been extensively studied in several species, including *Oryza sativa* (Xie *et al*., 2012)*, Nicotiana tabacum* (Feng *et al*., 2016), *Pyrus pyrifolia* (Qian *et al*., 2017), *Solanum lycopersicum* (Silva *et al*., 2014), *Passiflora edulis* (Silva *et al*., 2019), and *Populus tremula* (Lawrence *et al*., 2021). The miR156/SPL module is associated with leaf maturation, trichome formation, male fertility (anther development), phytomer length and composition, glucose and trehalose metabolism, as well as stress responses (Wang *et al*., 2009; Xing *et al*., 2010; Yu *et al*., 2010; Poethig, 2013; Fouracre & Poethig, 2016; Silva *et al*., 2019; Lawrence *et al*., 2020). Interestingly, several miRNA regulatory modules have been shown to be important for controlling the biosynthesis of secondary metabolites, including the miR156/SPL module (Padhan *et al*., 2016; Samad *et al*., 2020). At present, few examples connect the miR156-targeted and non-targeted *SPLs* with the control of the secondary metabolism. MiR156-targeted *SPL9* directly binds to the promoter of the sesquiterpene synthase gene *TPS21* to activate terpene synthesis (Yu *et al*., 2014). In *Arabidopsis*, the miR156-targeted *SPL9* negatively regulates anthocyanin accumulation by directly preventing expression of anthocyanin biosynthetic genes (Gou *et al*., 2011). *SPLs* may also directly or indirectly influence the carotenoid formation in plants. The tomato colorless nonripening (*Cnr*) locus encodes an *Arabidopsis* miR156-targeted *SPL3* homologue, and the *Cnr* mutation leads to low concentrations of total carotenoids, resulting in tomato fruits with a white pericarp (Fraser *et al*., 2001; Manning *et al*., 2006). More recently, Banana *(Musa acuminata) MuSPL16* was shown to be a positive regulator of carotenoid production through direct activation of carotenoid biosynthetic genes (Zhu *et al*., 2020), whereas papaya *(Carica papaya) SPL1/SBP1* is a direct transcriptional repressor of the *PHYTOENE DESATURASE4 (PDS4)*, which encodes a key enzyme involved in the conversion of phytoene into lycopene (Han *et al*., 2019).

Although the control of the biosynthetic pathway of apocarotenoids in *B. orellana* is still poorly understood, recent evidences indicated that bixin biosynthesis is modulated by developmental cues. In seeds, for instance, bixin levels increased during maturation (Moreira *et al*., 2022). We hypothesized that the miR156/SPL module regulates bixin production in annatto leaves in an age-dependent manner. To test this conjecture, we previously generated transgenic annatto plants overexpressing the miR156 (Faria *et al*., 2022). Here, we showed that the loss of miR156-targeted *SPL* function prolonged the juvenile phase in annatto, leading to reduced bixin levels in leaves as late as 90 days post-germination. Low expression of several *SPL* genes in miR156-overexpressing annatto plants impacted leaf ontogeny, modifying the expression patterns and accumulation of the main enzymes involved in bixin and ABA biosynthetic pathways. Together, our findings suggested that the carbon flux in miR156-overexpressing *B. orellana* leaves was somewhat redirected from bixin to ABA production, indicating an age-dependent leaf dynamic of bixin biosynthesis.

## Materials and Methods

### Plant material and growth conditions

Two OE::156 lines of *B. orellana* cv. Piave Vermelha were selected based on miR156 expression levels. Eight cloned plants of each OE::156 line and four No-transformed (Nt) lines at T0 generation were acclimatized as described by Faria *et al*. (2019). Plants were transferred to 12-L polyethylene pots with horticultural soil conditioner substrate (Tropstrato HT Hortaliças.Vida Verde Indústria e Comércio de Insumos Orgânicos Ltda, Mogi Mirim, SP, Brazil), and maintained under greenhouse conditions. The fertilizer Osmocote®Plus 16-08-12 5-6M (ICL, USA) was added (2 g per pot) on a bimonthly basis. The plants were watered daily and maintained at an 11/13-h light/dark photoperiod and 25/16 °C (day/night). Access to genetic heritage adhered strictly to current Brazilian biodiversity legislation and was approved by the Brazilian National System for the Management of Genetic Heritage and Associated Traditional Knowledge (SISGEN) under permission number AF3BC47. All GMO manipulations were performed in the Laboratório de Cultura de Tecidos II – Bioagro and in a GMO-adapted greenhouse following the CTNBio and CIBio-UFV biosafety rules.

### Phenotyping and microscopy

Acclimatized OE::156 lines were maintained under greenhouse conditions and evaluated at 15, 30, 45, 60, 75, and 90 days after planting (DAP). The following variables were assessed: total leaf number, plant height (cm), number of phytomers, diameter of the main axis, length of the third leaf (from petiole insertion to the apical portion of the leaf blade), leaf area, and ramification index (total ramification length and main plant axis length) (Morris *et al*. 2001). We measured the area of all Nt leaves and then sampled the leaf area of OE::156 plants using the same number of leaves as in Nt plants. The total OE::156 leaf area was estimated as sampled leaf area × 15, because OE::156 plants had 15 times more leaves than Nt plants. All images were analyzed using ImageJ software version 1.43u (National Institutes of Health, Bethesda, MD, USA) (Schneider *et al*., 2012).

For scanning electron microscopy analysis, stems of Nt and OE::156 were cut longitudinally in the third phytomer rootwards, because the extrafloral nectaries (EFN) were secreting in Nt plants. Then, the sectioned fragments were transferred to Karnovsky solution 0.1 M (Karnovsky, 1965) under −250 mm Hg vacuum for 1 h. The samples were dehydrated in a crescent ethanol series (50% to 100%) under −250 mm Hg vacuum in each dehydration step. Following, the samples were critical point dried with CO2 (CPD 030, Balzers), and metallized with a thin layer of gold (20 nm) in a sputter-coater (Quorum Q150RS). Samples were examined in a scanning electron microscopy (LEO 1430 VP) at an acceleration voltage of 10.6 kV.

### RNA extraction and cDNA synthesis

Expression profiles of bixin- and carotenoid-related genes, miR156, miR172, and *SPLs* in annatto were assessed by quantitative real-time PCR (qRT-PCR) using the CFX96 Touch™ Real-Time PCR Detection System (Bio-Rad, Hercules, CA, USA). Approximately 10 mg of freeze-dried third fully expanded leaves from 3-months-old plants were ground in liquid nitrogen and used for this analyzes. The final reaction volume of 10 μL included 40 ng cDNA, 400 nM primers, SYBR-Green Supermix (Bio-Rad), and deionized water. Expression data were derived from four biological replicates, with at least two technical replicates. Sequences of primers used in this study are listed in Table S1. Expression data were normalized using the *40S RIBOSOMAL PROTEIN S9 (BoRPS9)* (Moreira *et al*., 2018; Faria *et al*., 2022) (Table S1), and expression levels of each gene and miRNA were calculated using the 2^−ΔΔCt^ method (Livak and Schmittgen, 2001). Differences were compared using Student’s *t*-test (*P* ≤ 0.05).

### Identification, annotation, gene structure, and phylogenetic analysis of *BoSPL* gene family

To identify annatto sequences containing SBP/SPL domain (PF03110), all 16 *Arabidopsis (A. thaliana)* and 19 rice *SPL* gene family (Cardon *et al*., 1999; Xie *et al*., 2006) were downloaded from TAIR (https://www.arabidopsis.org/) and Phytozome v13 (https://phytozome-next.jgi.doe.gov/) and used as a query to conduct local BLASTn against all 73.381 transcripts generated by RNA-Seq from *Bixa orellana* (Moreira *et al*., 2022). All retrieved CDS were translated in all six frames using EMBOSS Transeq (https://www.ebi.ac.uk/Tools/st/emboss_transeq/) and analyzed by PFAM v35.0 (Salazar *et al*., 2020) to maintain only those containing the SBP/SPL domain (Table S2). The coding region of each *BoSPL* gene was predicted through gene structure display server (*GSDS* 2.0) program (http://gsds.gao-lab.org/, Hu *et al*., 2014).

All unique and primary coding sequences from *S. lycopersicum* (ITAG4.0 assembly), *T. cacao* (v2.1 assembly), *P. trichocarpa* (v4.1 assembly), *V. vinifera* (v2.1 assembly), and *P. virgatum* (v4.1 assembly) were downloaded at Phytozome v13 and used to recovery sequences containing PF03110 domain. The *SBP* or *SPL* gene names were used as described previously for tomato (Salinas *et al*., 2012), poplar (Guo *et al*., 2021; Hou *et al*., 2013), grapes (Hou *et al*., 2013; Díaz-Riquelme *et al*., 2012), and switchgrass (Wu *et al*., 2016).

All protein sequences for each multigene family were aligned using the default settings of <u>Mu</u>ltiple <u>S</u>equence <u>C</u>omparison by <u>L</u>og-<u>E</u>xpectation (MUSCLE) (http://www.ebi.ac.uk/Tools/msa/muscle/; Madeira *et al*., 2022). A Maximum Likelihood (ML) phylogenetic analysis was performed as using PhyML3.0 (Guindon *et al*., 2010), using Smart Model Selection (SMS) to select the best model (Lefort *et al*., 2017) (like VT+R+F substitution model) and aLRT branch support testing (Anisimova & Gascuel, 2006). The phylogenetic tree was visualized using the iTOL software (https://itol.embl.de; Letunic & Bork, 2021).

### miR156 and miR172 identification, and target *SPL* prediction in *Bixa orellana*

To identify miR156 and miR172 mature sequences, microRNA database available for *Arabidopsis*, tomato, cocoa, grape, poplar, and rice were used, such as *miRBase* (http://www.mirbase.org/search.shtml, Kozomara & Griffiths-Jones, 2010) and *PMRD: plant microRNA database* (http://bioinformatics.cau.edu.cn/PMRD/; Zhang *et al*., 2009). The identification of miR156 target regions *in silico* and *SPLs* genes sequences in *Bixa orellana* was performed through multiple alignments in *MUSCLE* and *psRNATarget* (https://www.zhaolab.org/psRNATarget/; Dai *et al*., 2018), using 20-21 nt miR156 sequences.

### Bixin content and bixin channel quantification

Bixin content was quantified as described (Napoleão *et al*., 2017; Faria *et al*., 2019). Approximately 10 mg of freeze-dried third fully expanded leaves from 3-months-old plants were ground in liquid nitrogen, followed by repeated extraction with 300 μL methanol:isopropanol:acetic acid (20:79:1; v/v/v). Samples were agitated four times for 20 s, sonicated for 10 min, kept on ice for 30 min, and centrifuged at 20,000 × *g* for 10 min at 4 °C. The supernatant was collected, filtered (Econofltr PVDF 13 mm, 0.2 μm; Agilent Technologies, Santa Clara, CA, USA), and analyzed by liquid chromatography-tandem mass spectrometry (LC-MS/MS). Samples of the final supernatant (5 μL) were automatically injected into a triple quadrupole (QqQ) LC-MS instrument (6430; Agilent Technologies, Waldbronn, Germany) equipped with an Eclipse Plus C18 column (2.1 × 50 mm, 1.8 μm) and a Zorbax SB-C18 guard column (1.8 μm, Agilent Technologies) at 26 °C. The mobile phase consisted of 0.02% acetic acid in water (solvent A) and 0.02% acetic acid in acetonitrile (solvent B), at a constant flow rate of 300 μL min^−1^. A linear gradient was applied as follows: 0–3 min, 2% to 60% B; 3–8 min, 60% to 99% B; 8–11 min, 99% B; 11–12 min, 99% to 2% B; and 12–15 min, 2% B. An electrospray ionization source was used under the following conditions: gas temperature of 300 °C, nitrogen flow rate of 10 L min^−1^, nebulizer pressure of 35 psi, and capillary voltage of 4,000 V. Bixin was analyzed via multiple reaction monitoring of ion pairs with mass transitions (395/157) and quantified via calibration curves using pure standards (1–200 μg and 0.1–500 ng; Sigma-Aldrich, St. Louis, MO, USA). Data were analyzed using Mass Hunter Workstation software (Agilent Technologies).

For bixin channels counting, images of the abaxial leaf surfaces of 3-month-old Nt, OE::156_1, and OE::156_4 plants were captured using a stereomicroscope (SZX7; Olympus, Tokyo, Japan) coupled to a Moticam 580 5.0 MP camera (Nikon, Tokyo, Japan). Images were analyzed using ImageJ version 1.49 (Schneider *et al*., 2012). Bixin channels were quantified by sampling the basal, medial, and apical leaf portions, with three different areas per image, using three leaves per genotype. The number of bixin channels was divided per mm^2^. Differences were compared using Student’s *t*-test (*P* ≤ 0.05).

### Abscisic acid (ABA) profile

ABA levels were quantified in the eight third (from apex to base) fully expanded leaves collected from 3-month-old greenhouse-grown Nt and OE::156 plants. Leaf samples (approximately 110 mg) were powdered in liquid nitrogen. A 300-μL aliquot of extracting solution (methanol:isopropanol:acetic acid 20:79:1) was added to 1.5 mL microtubes. The samples were vortexed four times for 20 s each, sonicated for 5 min, kept at 4 °C for 30 min, and centrifuged at 13,000 × *g* for 10 min and 4 °C. Subsequently, 350 μL of supernatant was collected in a new microtube. The pellet was run through the same steps, and the supernatant was collected. ABA identification and quantification was carried out by LC-MS/MS using an Agilent 1200 Infinity Series chromatograph coupled to a 6430 triple quadrupole mass spectrometer as described by Napoleão *et al*. (2017). Differences were compared using Student’s *t*-test (*P* ≤ 0.05).

### Pigment quantification

Chlorophyll and carotenoid contents were quantified (μg mL^−1^) following the protocols described by Wellburn (1994) and Warren (2008). Briefly, 600 μL of 100% methanol was added to 50 mg of freshly frozen macerated leaf samples of 3-months-old plants. The solution was vortexed for 30 s and incubated at 4 °C with agitation in the dark for 10 min. The samples were centrifuged at 12,000 × *g* (4 °C, 10 min), and the supernatant was collected. The procedure was repeated for the remaining pellet, and the resulting supernatant was collected and mixed with the first one. After extraction, the supernatant (200 μL) was collected and added to a microplate for absorbance readings (A) at 470, 653, and 666 nm in a Multiskan FC microplate reader (Thermo Fisher Scientific, Waltham, MA, USA). Total chlorophyll (Chl *a* + Chl *b*) and carotenoid contents were determined according to the following equations:

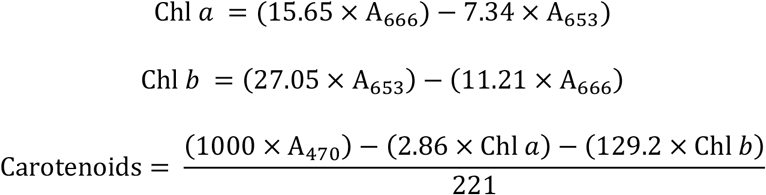

Anthocyanins were quantified as described by Neff and Chory (1998) using the same samples as for chlorophyll/carotenoid measurements, but at 657 and 530 nm. Relative anthocyanin content was expressed as (A_530_ - A_657_) mg^−1^ fresh weight. Differences were compared using Student’s *t*-test (*P* ≤ 0.05).

### Proteomic extraction and analysis

For proteomic quantification, leaf samples were collected from 3-month-old Nt and OE::156_4 plants. Total protein extraction was performed according to Damerval *et al*. (1986) with some modifications. Briefly, 10 mg of freeze-dried pulverized leaves in liquid nitrogen were mixed with 1 mL of chilled solution containing 10% (w/v) trichloroacetic acid (TCA; Sigma-Aldrich) in acetone (Merck; Darmstadt, Germany) and 20 mM DTT (GE Healthcare, Piscataway, USA), and the mixture was vortexed and centrifuged (Reis *et al*., 2021). The resulting pellets were washed three times with cold acetone containing 20 mM DTT and centrifuged for 5 min in each wash step. The pellets were air dried and resuspended in solubilization buffer containing 7 M urea, 2 M thiourea, 2% Triton X-100, 1% dithiothreitol (all GE Healthcare, Piscataway, NJ, USA), 1 mM phenylmethanesulfonyl fluoride (Sigma-Aldrich), and complete protease inhibitor cocktail (Roche Diagnostics, Mannheim, Germany). The mixture was vortexed and centrifuged. We then collected the supernatants and measured the protein.

The protein samples were precipitated using methanol/chloroform to remove any contaminants (Nanjo *et al*., 2012), resuspended in a solution consisting of 7 M urea and 2 M thiourea, and digested using Microcon-30 kDa filter units (Millipore, Billerica, MA, USA) according to the filter-aided sample preparation methodology described by (Reis *et al*., 2021). After digestion, the resulting peptides were quantified via absorbance measurements at 205 nm using a NanoDrop 2000c spectrophotometer (Thermo Fisher Scientific) and then injected into a NanoAcquity ultra-performance LC coupled to a Q-TOF SYNAPT G2-Si mass spectrometer (Waters, Manchester, UK).

Spectral processing and database searches were performed using Protein Lynx Global SERVER software version 3.02 (Waters), and label-free quantification analyses were performed using ISO Quant software version 1.7 (Distler *et al*., 2014). Protein identification was performed against a nonredundant protein databank for *B. orellana* generated by transcriptome sequencing and *de novo* assembly (Moreira *et al*., 2022). The mass spectrometry proteomics data have been deposited to the ProteomeXchange Consortium via the PRIDE (Perez-Riverol *et al*., 2022) partner repository with the dataset identifier PXD036944. Only proteins that were present or absent (for unique proteins) in all three runs of biological replicates were considered in the differential accumulation analysis. Proteins with significant Student’s t test (two-tailed; *P* < 0.05). Proteins with significant Student’s *t*-test (two-tailed, *P* ≤ 0.05) results were considered differentially accumulated (DAP), as up-regulated if the Log2 fold change (FC) was greater than 0.6 and down-accumulated if the Log2 FC was less than −0.6. For functional annotations, we used OmicsBox software (https://www.biobam.com/omicsbox) and UniProtKB (www.uniprot.org). Kyoto Encyclopedia of Genes and Genomes (KEGG) enrichment pathway analysis between DAPs (*P* ≤ 0.01) was performed in Metascape (Zhou *et al*., 2019) after a BLAST search of NCBI (https://www.ncbi.nlm.nih.gov) to obtain *Arabidopsis* reference sequences. The predicted protein interaction networks were constructed using *A. thaliana* homologs in *B. orellana*, which were identified through a STRING search followed by downstream analysis in Cytoscape version 3.9.1 (Shannon *et al*., 2003).

### Statistical analysis

The software IBM®SPSS®v. 22 was used for all statistical analyzes. For outlier’s detection and removal, the boxplot method was used. In the phenotyping parameters, qRT-PCR analyzes, pigment content, bixin and ABA measure, and bixin channels number, the means differences of each OE::156 line were compared with the control Nt, using Student’s *t*-test (*P* ≤ 0.05).

## Results

### Low levels of miR156-targeted *SPLs* affect growth and leaf shape of *B. orellana*

We have previously generated *B. orellana* plants overexpressing the miR156 (Faria *et al*., 2022). For phenotypic analyses, we selected two *in vitro* OE::156 lines with similar vegetative architecture and ramification patterns to avoid any bias that would affect subsequent growth *ex vitro*. Moreover, flow cytometric analyses of these two lines showed stability of nuclear DNA compared with non-transgenic lines (Faria *et al*., 2022). To our knowledge, to date there are no studies characterizing the transition between juvenile and adult phases in *B. orellana* plants. Therefore, we initially compared vegetative architecture and leaf development between OE::156 and Nt plants growing under greenhouse conditions. In general, three-month-old OE:: 156 plants displayed drastically modifications in canopy growth patterns, with multiple shoots arising from axillary buds (Fig. 1a). In addition, OE::156 plants had smaller leaves compared to their Nt counterparts (Fig. 1b). These observations suggested that larger heart-shaped leaves may be a morphological marker for phase change in *B. orellana*.

**Figure 1.**
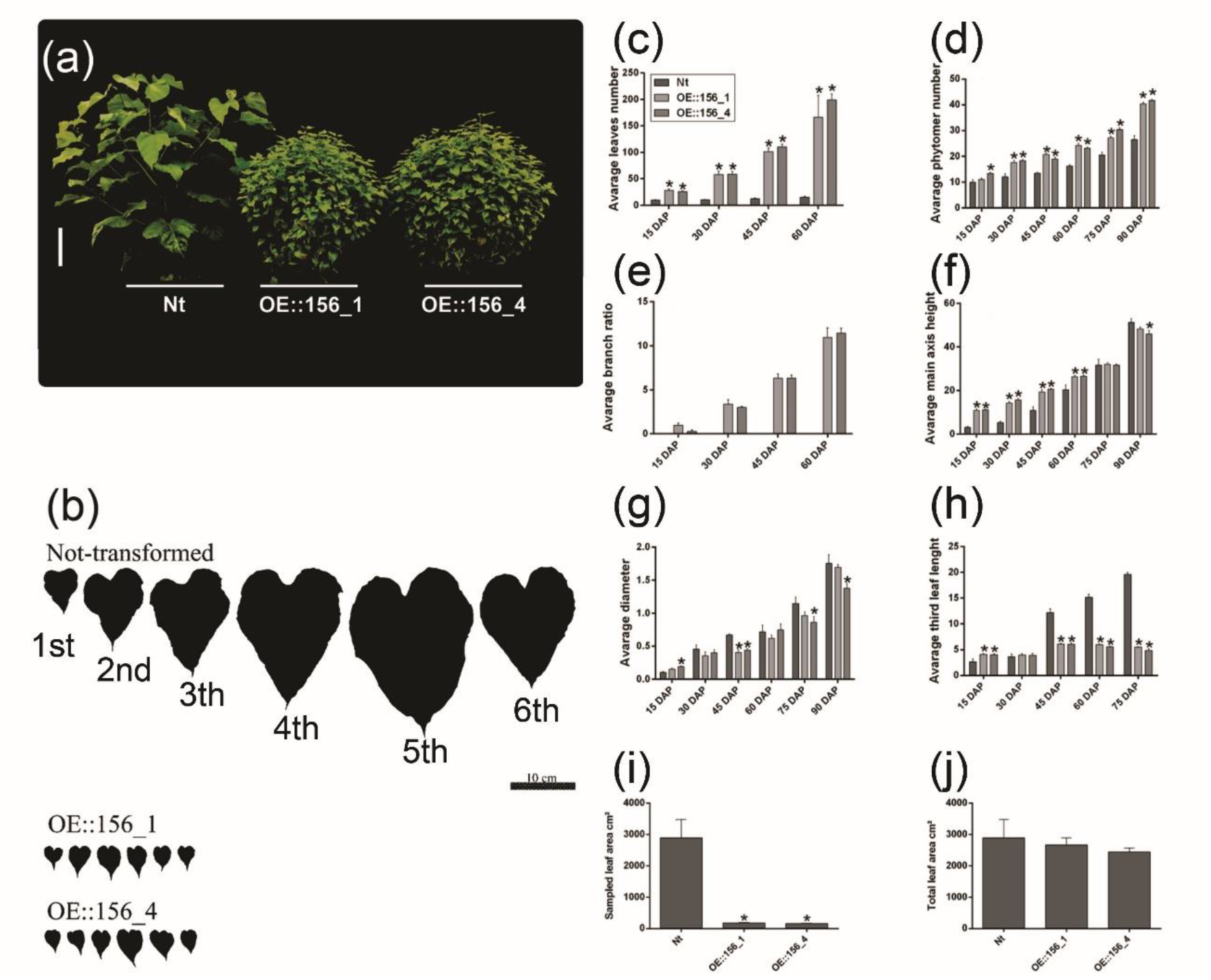
miR156-targeted *SPLs* are required for proper canopy architecture and leaf development in *B. orellana*. **(a)** Vegetative architecture and size of non-transformed (Nt) plants and plants overexpressing miR156 (OE::156) in two distinct lines (OE::156_1 and OE::156_4) on the 90^th^ day after planting in a greenhouse. Bar: 10 cm. **(b)** Leaf shape and size of Nt and OE::156_1 and OE::156_4 lines. Bar: 10 cm. From left to right: first to sixth leaf **(c)** Total leaf number until 60 days after planting (DAP) in Nt, OE:: 156_1, and OE:: 156_4 plants. **(d)** Average phytomer number assessed up to 90 DAP in Nt, OE::156_1, and OE::156_4 plants. **(e)** Branching index assessed up to 60 DAP in Nt, OE::156_1, and OE::156_4 plants. **(f)** Average values for main stem height assessed up to 90 DAP in Nt, OE::156_1, and OE::156_4 plants. **(g)** Average values for main stem diameter until 90 DAP in Nt, OE:: 156_1, and OE::156_4 plants. **(h)** Third leaf length (from petiole insertion to the end of the leaf blade) 75 DAP in Nt, OE::156_1, and OE::156_4 plants. **(i)** Leaf area on the 90^th^ DAP (n = 20) in Nt, OE::156_1, and OE::156_4 plants. **(j)** Total leaf area [leaf area × number of Nt leaves (15)] on the 90^th^ DAP in Nt, OE::156_1, and OE::156_4 plants. \**P* ≤ 0.05 (Student’s *t*-test). Vertical bars denote the standard error.

Next, we characterized in detail greenhouse-grown plants every 15 days until the 90th day. As expected, both OE::156 lines produced more leaves than their Nt counterparts in all evaluated timepoints (Fig. 1c), as well as a higher phytomer number after 30 days of planting (Fig. 1d). Quantification of the branching ratio revealed 11-fold more branches in OE::156 plants than in Nt plants (Fig. 1e), indicating a strong loss of apical dominance. Stem height was higher in OE::156 plants than in Nt plants until the 75th day after planting, although the opposite was observed on the 90th day (Fig. 1f). Although not statistically significant in all timepoints evaluated, our data suggest that OE::156 plants displayed less secondary growth compared with Nt plants, as indicated by their reduced main stem diameter compared with Nt plants (Fig. 1g). We also evaluated leaf growth over time in Nt and OE::156 plants, analyzing the first fully expanded leaf of *B. orellana*. On the 15th day, OE::156 third leaves grew faster than Nt, but the opposite was observed from the 45th day on (Fig. 1h). Finally, we quantified leaf area by photographing the leaves after the 90th day. Because of the high number of leaves in OE::156 plants (Fig. 1a-c), we used an equivalent number of leaves in OE::156 and Nt plants. Therefore, the leaf area for both genotypes was comparable. OE:: 156 leaf area was 16 times smaller than Nt plants (Fig. 1i). No significant differences were observed between total OE::156 and Nt leaf areas when the former was adjusted by multiplying the measured OE::156 leaf area, which is the same amount of Nt leaves, by 15, because OE leaves have 15 times more leaves than Nt plants (Fig. 1j). These findings implied that OE::156 lines produced more leaves with distinct ontogenetic growth, but with overall leaf area index similar to Nt plants, indicating that the loss of *SPL* function delayed phase transition in *B. orellana*.

Leaf shape is commonly used to distinguish juvenile versus adult identity, but light intensity can sometimes affect leaf shape without necessarily operating via age-dependent mechanisms (Jones, 1995). Therefore, we used additional traits to evaluate the apparent delay in phase changing of OE::156 lines. The presence of sugar-secreting extrafloral nectaries (EFN) is age-dependent in some trees. For instance, the appearance of EFNs produced by acacias (genus *Vachellia*) is tightly correlated with the decline in the expression of miR156 and increasing expression of *SPLs*, suggesting that these structures evolved by co-opting a preexisting age-dependent program (Leichty and Poethig, 2019). While Nt plants produced EFNs in several nodes, miR156-overexpressing *B. orellana* plants did not produce any visible EFNs and displayed apparently less trichomes in the stems (Fig. S1, S2), reinforcing that the miR156/SPL module coordinates the timing of vegetative development in achiote trees.

**Figure S1.**
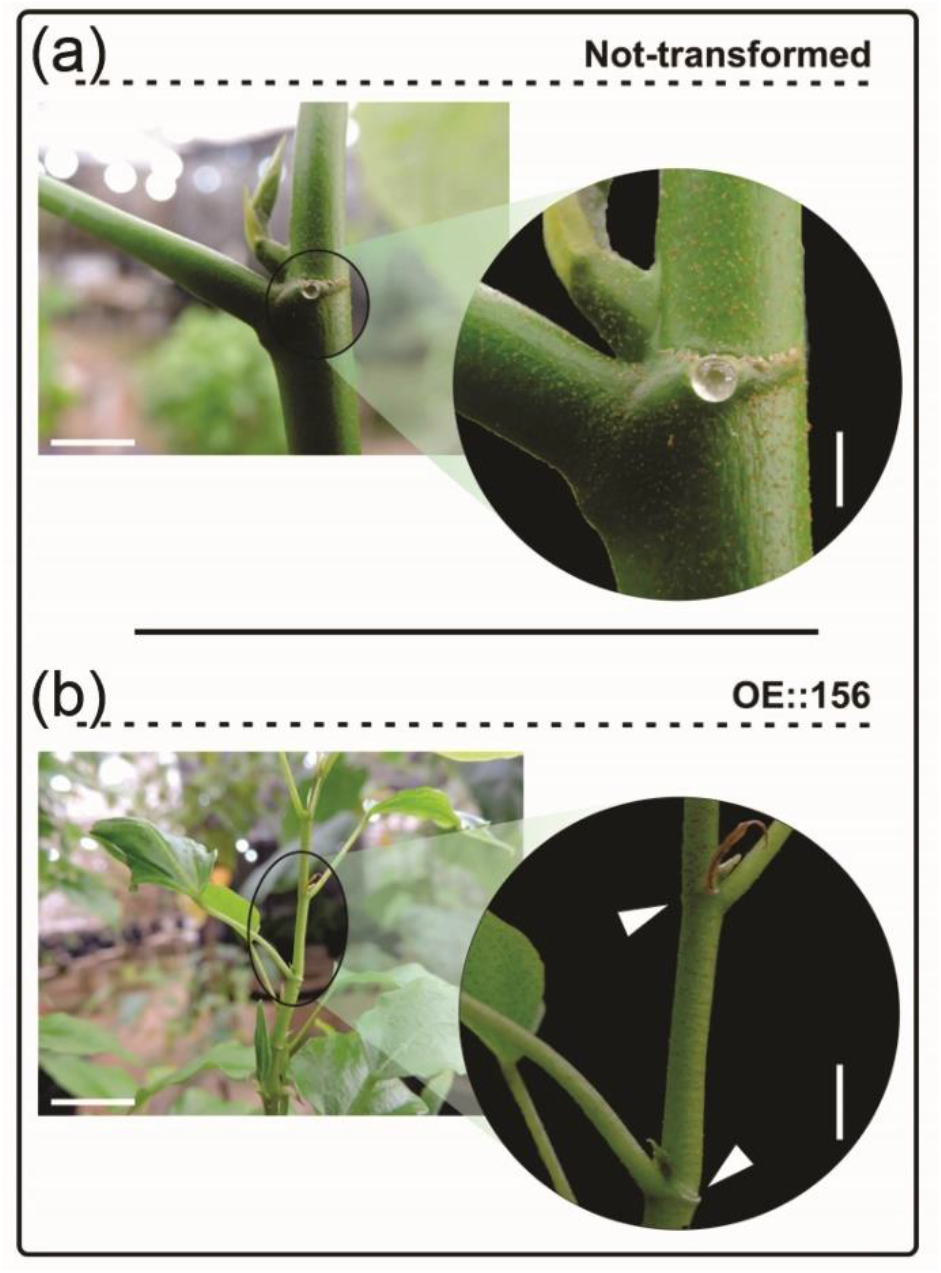
miR156-targeted *SPLs* are required for extrafloral nectary (EFN) formation. **(a)** Detail of EFNs exudating nectar in non-transformed (Nt) plants. **(b)** Detail of OE::156_4 plants, in which the total absence of EFN is indicated by arrows. Bars: 0.6 cm.

**Figure S2.**
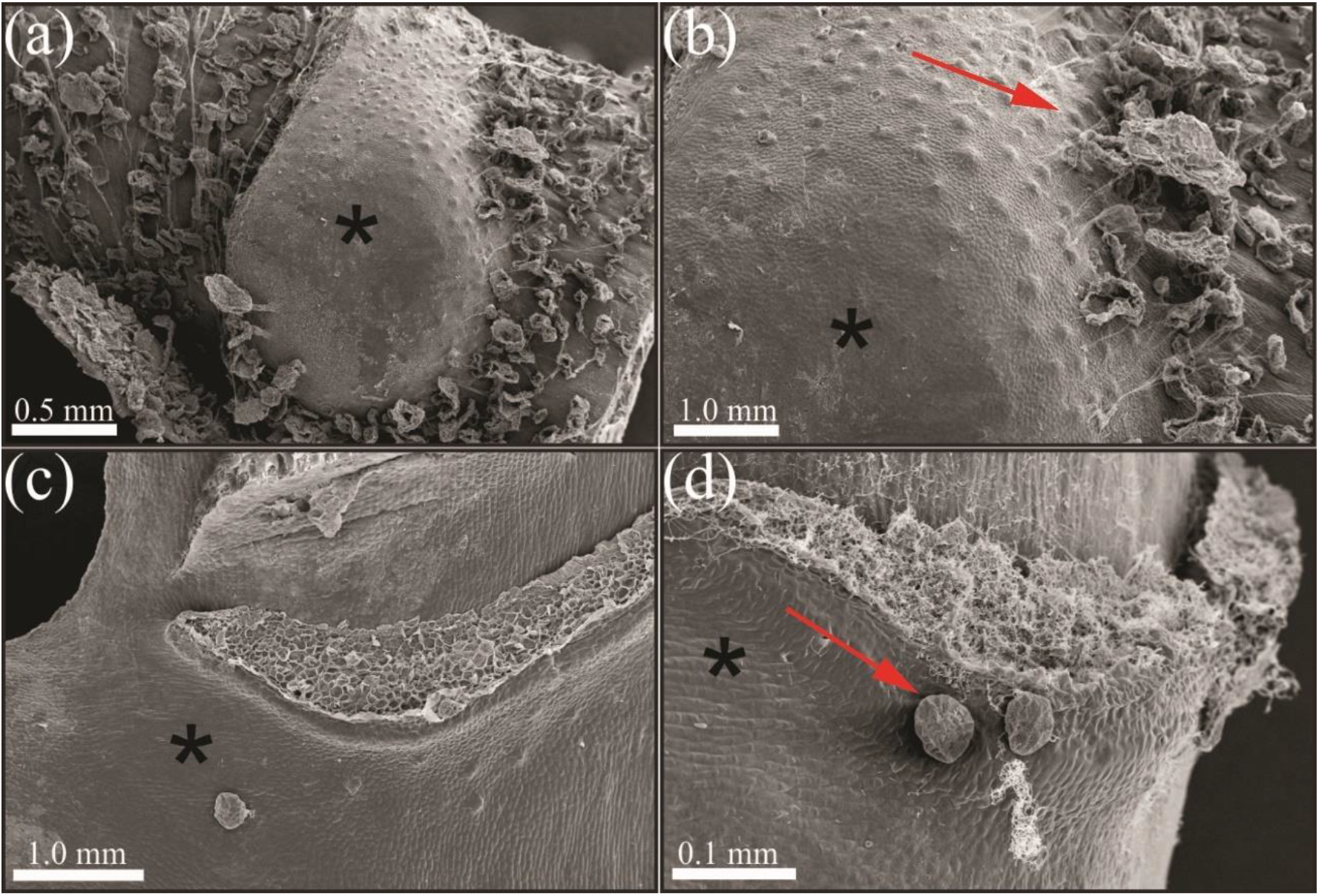
miR156-targeted *SPLs* are required for extrafloral nectary (EFN) and trichome formation. **(a)** EFNs in Non-transformed (Nt) stem. The asterisk indicates the secretory structure. **(b)** Trichomes (red arrow) in an EFN in Nt stem. The asterisk indicates the secretory structure. **(c)** OE::156 stem without presence of visible EFNs. The asterisk indicates where the secretory structure would be positioned in analogy with Nt. **(d)** Trichomes (red arrow) in a OE::156 stem. The asterisk indicates where the secretory structure would be positioned in analogy with Nt.

### MiR156-overexpressing leaves have reduced bixin content

Given that leaf development and ontogeny in *SPL*-defective *B. orellana* plants are distinct from Nt plants (Fig. 1a; S1; S2), we investigated whether such alterations in leaf growth affected bixin production. MiR156-overexpressing leaves displayed 2.5-fold less bixin levels than Nt (Fig. 2a), albeit no significant differences were observed in the overall pigment content (Table S3). We then hypothesized that modifications in leaf structures, such as secretory structures, would be specifically impacted by *SPL* low levels in OE::156 plants. Bixin is produced in all plant tissues and accumulated in secretory structures, known as bixin channels (Almeida *et al*., 2021). Therefore, we quantified bixin channels on the abaxial face of the third leaf (Fig. 2b, c). In the basal region of the leaf, the number of bixin channels did not differ between OE::156 plants and Nt plants. In contrast, there were fewer bixin channels in the medial and apical parts of OE::156 leaves (Fig. 2c), which may be another fundamental adult trait of *B. orellana* as shown for *Arabidopsis* (Kang and Dengler, 2004). Given that the formation of bixin channels require structural maturation of these tissues (Almeida *et al*.,2021), as noted also by the differential leaf venation pattern (Fig. 2c; gray arrows), these findings suggested a negative correlation between leaf maturation and the loss of miR156-targeted *SPL* activity.

**Figure 2.**
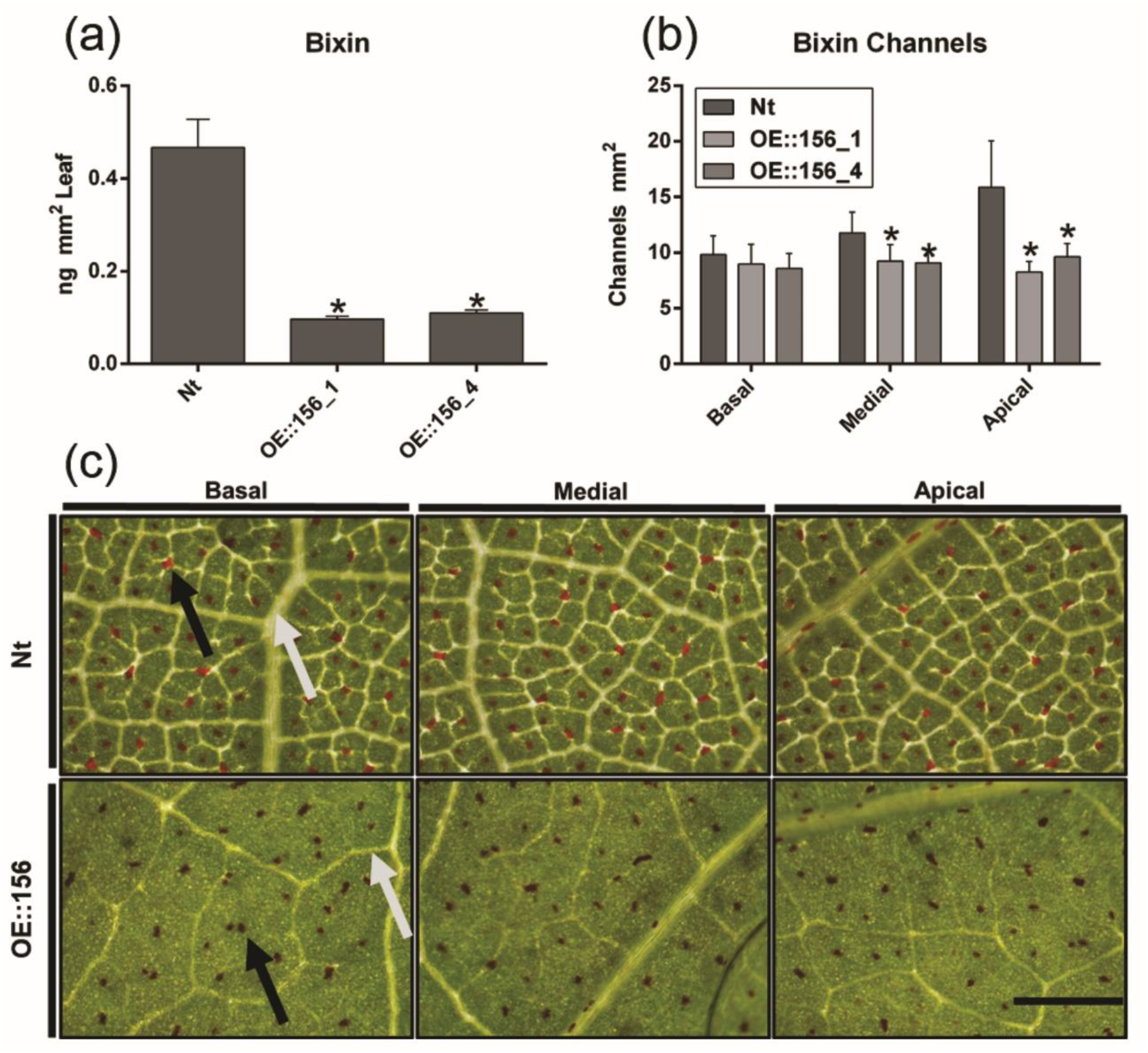
miR156-targeted *SPLs* modulate bixin-related features in *B. orellana*. **(a)** Average total bixin content in *B. orellana* leaves from non-transformed (Nt) plants and plants overexpressing miR156 (OE::156_1 and OE::156_4) on the 90^th^ day after planting (DAP). The results are adjusted to the mass of fresh leaves from four plants of each genotype. **(b)** Average number of bixin channels in different *B. orellana* abaxial leaf parts on the 180^th^ DAP. **(c)** Detail of bixin channels in *B. orellana* leaves on the 180^th^ DAP. The black arrows point to bixin channels and the gray arrows indicate veins on the abaxial face of the leaf blade. Bar: 1 cm. \**P* ≤ 0.05 (Student’s *t*-test); vertical bars denote the standard error.

### Identification, phylogenetic analysis, and expression patterns of miR156, miR172 and *BoSPLs*

To identify miR156-targeted *SPL* genes, we took advantage of the recently published *B. orellana* transcriptomes in several organs, including leaves and seeds (Cárdenas-Conejo *et al*., 2015; Moreira *et al*, 2022). Our data revealed a set of 13 annatto *SPLs* family members, designated as *BoSPL* and their respective number considering its closest related phylogenetic *SPLs*/*SBPs* gene, following *Arabidopsis*, cacao, and grapes, respectively (Fig. 3 and Table S2). With exception of VIII clade, all 13 *BoSPLs* genes were found into eight known clades (Salinas *et al*., 2012). From the five predicted miR156 target clades (Fig. 3), five annatto genes (*BoSPL2, 6, 10, 13a*, and *13b*) showed recognition site for miR156 (Fig. 3f; Fig. S3), while four (*BoSPL3, 4, 5a*, and *5b*) did not (Table S4), probably due to the incomplete transcript sequence available.

**Figure 3.**
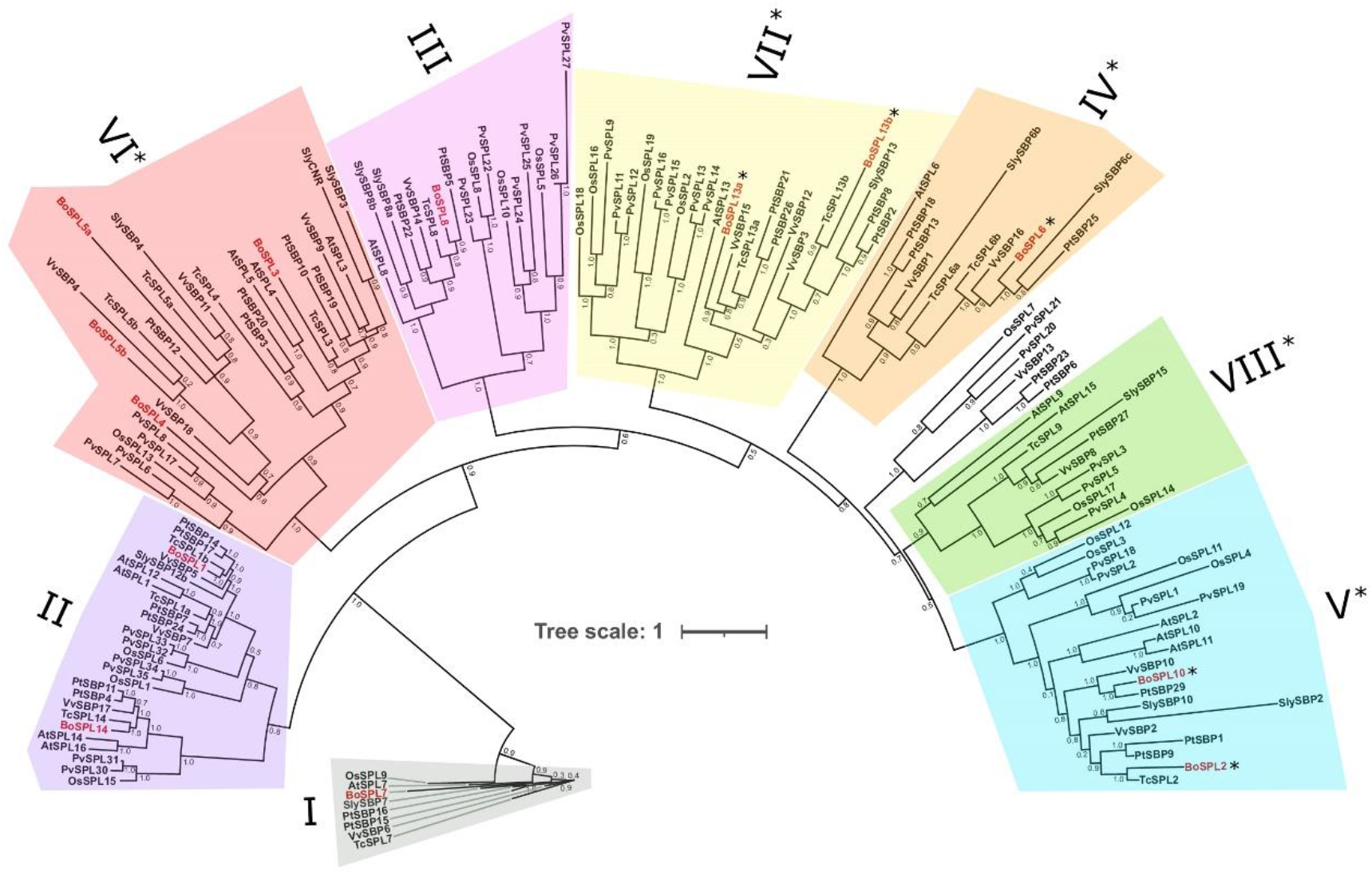
Maximum-likelihood phylogeny of the SPL/SBP proteins of *Bixa orellana* among *Arabidopsis*, tomato, cacao, grapes, rice, and swithgrass. Unrooted phylogenetic tree based on an alignment of all SPL/SBP sequence encoded protein obtained for *B. orellana (BoSPL)*, tomato (*SlySBP*), Arabidopsis (*AtSPL*), poplar (*PtSBP*), grapes (*VvSBP*), rice (*OsSPL*), and swithgrass *(PvSPL)*, using MUSCLE tool and Interative Tree of Life (iTOL) resource to annotate. Original code gene for each species is available (Table S2). Clade colors match Salinas *et al*. (2012) where applicable. Clades with asterisks show *SPLs/SBPs* regulated by miRNA156 or miR157, as stated by Preston and Hileman (2013). The code name of annatto *SPLs* were marked in red for highlight; those with asterisks contain recognition site for miR156 in their sequence. Scale bars indicate the number of substitutions per site.

**Figure S3.**
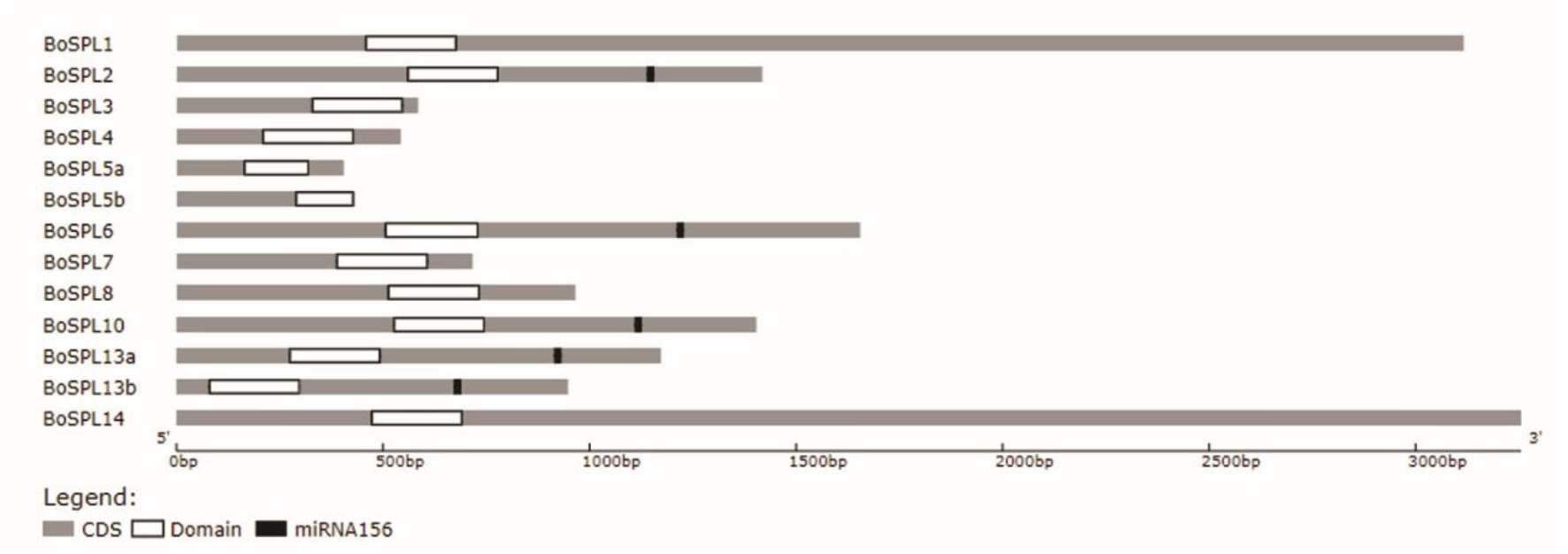
*BoSPLs* targeted by miR156. Light grey lines represent ORFs. The white box represents the conserved SBP domain. Prediction of miR156 recognition site in *BoSPLs* transcripts are shown in black box. Sequences were sized based on their nucleotide number (below).

To determine whether the observed *B. orellana* leaf phenotypes were consistent with high levels of miR156, we quantified its transcript levels by qRT-PCR in the greenhouse-grown plants. As expected, mature BomiR156 transcript abundance was significantly higher in both OE::156 lines compared with Nt (Fig. 4a), whereas the opposite was observed for BomiR172 (Fig. 4b). This observation is consistent with the opposing miR156 and miR172 expression patterns observed during *Arabidopsis* floral transition (Wang *et al*., 2009). We recently showed that miR172 transcript levels significantly increased in adult leaves of *Passiflora edulis*, whereas an opposite expression pattern was observed for miR156 (Silva *et al*., 2019). Collectively, these observations implied that miR156 and miR172 may have opposing roles associated with the leaf ontogenetic process in both *B. orellana*. We then selected three miR156-targeted *BoSPLs* (*6*, *13a*, and *13b;* asterisks at Fig. 3) to evaluate their transcript levels in OE::156 leaves. PCR primers were designed to a unique sequence within the predicted coding region of each *BoSPL* gene showing the miR156 recognition site (Fig. S3). These primers were then used to measure the abundance of these transcripts in OE::156 and Nt leaves. All three *BoSPLs* were repressed in OE:: 156 lines (Fig. 4c-e), suggesting that they might be associated with the distinct leaf ontogeny and low bixin levels observed in the miR156-overexpressing annatto leaves.

**Figure 4.**
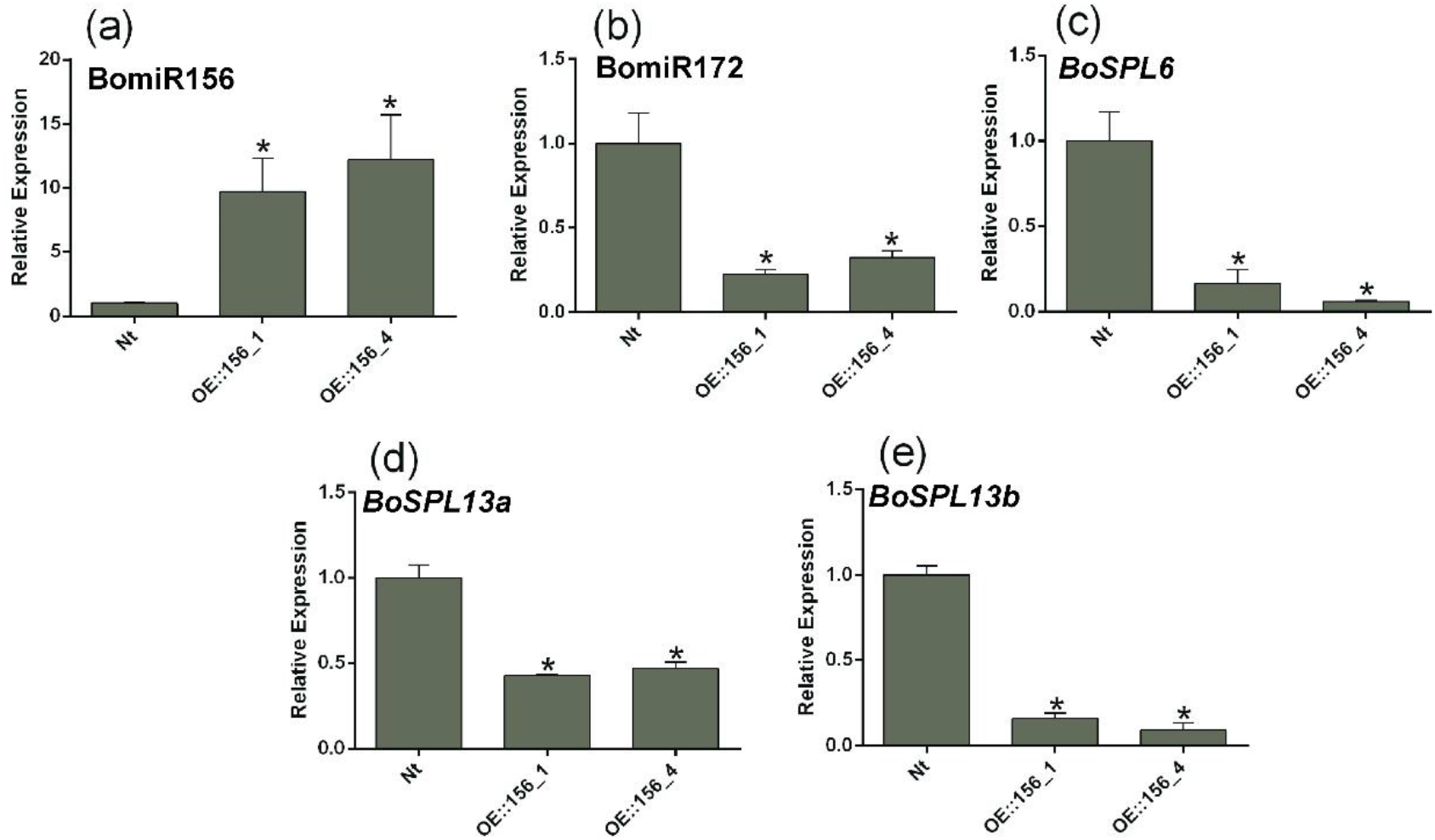
Highly expression of miR156 leads to low levels of miR172 and *BoSPL* transcripts in *B. orellana* leaves. Average expression profiles by qRT-PCR (relative expression) of *B. orellana* miR156 (BomiR156) (**a**), BomiR172 **(b)**, *BoSPL6* **(c)**, *BoSPL13a* **(d)**, and *BoSPL13b* **(e)** in leaves from non-transformed (Nt) plants and plants overexpressing miR156 (OE::156_1 and OE:: 156_4) on the 90^th^ day after planting in the greenhouse. \**P* ≤ 0.05 (Student’s t-test; n=4); vertical bars denote the standard error.

### MiR156-targeted *BoSPLs* modulate the expression of carotenoid and bixin biosynthetic genes

Transcriptomic profiling of distinct developmental *B. orellana* seed stages showed differential gene expression associated with carotenoid and bixin biosynthesis (Moreira *et al*., 2022). These results indicated that such genes are developmentally regulated, perhaps via the miR156/*SPL* module. To test this conjecture, we analyzed the expression of genes associated with carotenoid and bixin biosynthesis in OE::156 and Nt leaves. *B. orellana 1-DEOXY-D-XYLOSE-5-PHOSPHATE SYNTHASE 2A (BoDXS2a)* is responsible for converting pyruvate and glyceraldehyde-3-phosphate (GADP) into 1-deoxy-D-xylulose-5-phosphate, one of the first steps of lycopene (one of the bixin precursors) biosynthesis (Vaccaro *et al*., 2014). *BoDXS2a* was repressed in OE::156_1 compared with Nt leaves (Fig. 5a), but as OE::156_4 line did not differ from Nt, bixin production might be regulated downstream of *BoDXS2a*. Transcripts of annatto *PHYTOENE SYNTHASE 1 (BoPSY1)*, which converts dimethylallyl pyrophosphate into phytoene (Fu *et al*., 2014), were ~2.0-fold less abundant in OE::156 than in Nt leaves (Fig. 5b). This result suggested that the carotenoid biosynthesis pathway was generally suppressed in OE::156 plants.

**Figure 5.**
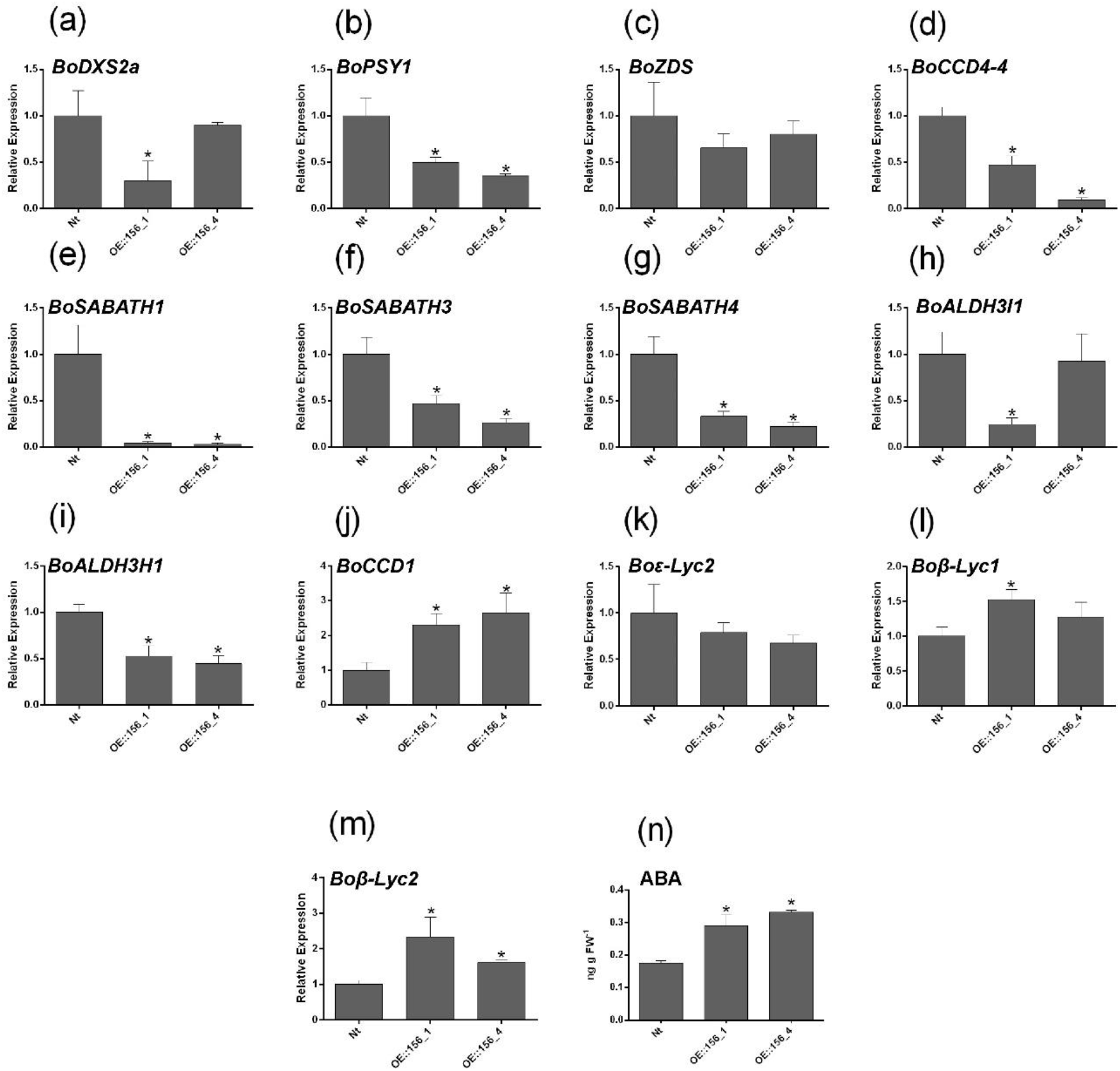
Expression levels of genes involved in carotenoid and bixin biosynthesis, as well as abscisic acid (ABA) accumulation in *B. orellana* leaves. *1-DEOXY-D-XYLOSE-5-PHOSPHATE SYNTHASE 2A (BoDXS2a)* **(a)**, *PHYTOENE SYNTHASE 1 (BoPSY1)* **(b)**, *ZETA-CAROTENE DESATURASE* (*BoZDS*) **(c)**, *CAROTENOID CLEAVAGE DIOXYGENASE4 (BoCCD4-4)* **(d)**, SABATH methyltransferase genes *(BoSABATH1)* **(e),** *BoSABATH3* **(f)** *BoSABATH4* **(g)**, *ALDEHYDE DEHYDROGENASE FAMILY 3 MEMBER I1 (BoALDH3I1)* **(h)**, *ALDEHYDE DEHYDROGENASE FAMILY 3 MEMBER H1 (BoALDH3H1)* **(i)**, *CAROTENOID CLEAVAGE DIOXYGENASE1 (BoCCD1)* **(j)**, *LYCOPENE ε-CYCLASE2 (Boε-Lyc2)* **(k)**, *LYCOPENE β-CYCLASE 1* (*Boβ-Lyc1*) **(l)** and *-2* (*Boβ-Lyc2*) **(m)**, and ABA **(n)** in non-transformed (Nt) plants and plants overexpressing miR156 (OE::156_1 and OE::156_4) on the 90^th^ day after planting in the greenhouse. \**P* ≤ 0.05 (Student’s *t*-test; n=4); vertical bars denote the standard error.

However, annatto *ZETA-CAROTENE DESATURASE (BoZDS)* mRNA levels did not significantly differ between OE::156 and Nt leaves (Fig. 5c). *CAROTENOID CLEAVAGE DIOXYGENASE4* (*BoCCD4-4*) is responsible for lycopene cleavage to generate bixin aldehyde, which is then fed into the bixin route (Carballo-Uicab *et al*., 2019). Among the *BoCCD4* genes, *BoCCD4-4* was expressed in leaves and throughout seed maturation (Moreira *et al*., 2022). Thus, we evaluated *BoCCD4-4* expression, and observed that *BoCCD4-4* mRNA levels were strongly reduced in OE:: 156 leaves (Fig. 5d), which indicates that miR156-targeted *SPL* function may be required for carotenoid production in *B. orellana* leaves. The SABATH methyltransferase gene family appears to methylate norbixin to bixin, and represents the likely end point of bixin production (Cárdenas-Conejo *et al*., 2015; Faria *et al*., 2020). *SABATH1, −3*, and *-4* are expressed in leaves (Moreira *et al*., 2022), and all three *SABATH* genes were down-regulated in OE::156 leaves (Fig. 5e, f, g). In particular, *BoSABATH1* expression showed an almost 8-fold reduction in both transgenic lines. Annatto *ALDEHYDE DEHYDROGENASE FAMILY 3 MEMBER I1* (*BoALDH3I1*) is also responsible for converting bixin aldehyde into norbixin (Cárdenas-Conejo *et al*., 2015). *BoALDH3H1-1* transcripts accumulated in leaves and seeds (Cárdenas-Conejo *et al*., 2015). Low levels of *BoALDH3H1-1* were observed in OE::156 compared with Nt leaves (Fig 5h). However, given that OE::156_4 plants did not differ from Nt, we tested another *BoALDH* gene, *BoALDH3H1* (Fig. 5i), which was repressed in both lines compared with Nt leaves.

Attempting to determine the fate of the carbons normally destined for bixin production, we initially quantified annatto *CAROTENOID CLEAVAGE DIOXYGENASE1* (*BoCCD1*) transcript levels. BoCCD1 may diverge the carbon flux away from bixin production in *B. orellana*(Cárdenas-Conejo *et al*., 2015). OE::156 leaves exhibited more than 2-fold *BoCCD1* transcripts than Nt counterparts (Fig. 5j). Carotenoid biosynthesis proceeds through two different pathways, depending on the activity of two enzymes: LYCOPENE ε-CYCLASE and LYCOPENE *β-* CYCLASE. LYCOPENE ε-CYCLASE constitutes a key control point for the continuation of lycopene to lutein or *β*-carotene biosynthesis (Cazzonelli & Pogson, 2010; Fu *et al*., 2019). In our analyses, *LYCOPENE ε-CYCLASE2 (Boε-Lyc2)* mRNA levels did not differ between OE::156 and Nt leaves, although OE::156 showed slightly fewer transcripts (Fig. 5k). In contrast, *B. orellana LYCOPENE β-CYCLASES 1* and *2* (*Boβ-Lyc1* and *Boβ-Lyc2*) showed higher expression in OE::156 leaves compared with Nt counterparts (Fig. 5l, m). Together, these observations suggested that low levels of *SPLs* interfered with *β*-carotene biosynthesis in annatto leaves. Given that *β*-carotene-derived xanthophylls serve as biosynthetic precursors of ABA (Nambara and Marion, 2005), we assessed ABA levels in Nt and OE::156 leaves. Both transgenic lines presented higher ABA levels than in Nt (Fig. 5n), indicating a possible deviation of carbon flux from bixin to ABA biosynthetic pathway.

### Overall secondary metabolism is affected in OE::156 leaves as revealed by proteomic analysis

Low levels of *SPL* transcripts in miR156-overexpressing alfafa plants modify the pool of proteins associated with primary and secondary metabolism (Arshad *et al*., 2020). To determine how the miR156-targeted *SPLs* modulate the production of secondary metabolite-related proteins/enzymes in *B. orellana* leaves, we employed proteomic analysis of the OE::156_4 leaves, and compared with Nt using LC-MS (Table S4). A total of 255 differentially altered proteins was detected, of which 89 were up-regulated and 159 were down-regulated in OE::156 leaves. Gene ontology (GO) analysis (Table S4) revealed that approximately 6% of the down-regulated proteins were associated with cell cycle (e.g., tubulin beta-1 chain). A similar trend was observed for other GO-related categories, such as cellular component biogenesis, developmental processes, and cellular component organization. Classifying differentially expressed proteins into their respective KEGG pathways (Fig. S4b) revealed that most proteins associated with sugar and nucleotide metabolisms, amino acid biosynthesis, secondary metabolite biogenesis, and ribosome biosynthesis were less abundant in OE::156 leaves (Fig. 6a). We used proteomic data to select for secondary metabolite-related proteins affected by the down-regulation of *BoSPLs*. Proteins associated with flavonoid, anthocyanin, and lignin biosynthesis were the most altered by the miR156/*SPL* module. A volcano plot analysis of the enzymes related to secondary metabolism (Table S5) highlighted five key candidates linked to pigments and other secondary metabolite pathways (Fig. 6b). Among the low abundant enzymes, we found ZEAXANTHIN EPOXIDASE (ZEP), which catalyzes the conversion of zeaxanthin to violaxanthin, and ABA biosynthesis (Hieber *et al*., 2000). Other enzymes associated with secondary metabolism were FLAVONOL SULFOTRANSFERASE-LIKE, PHENYLALANINE AMMONIA-LYASE, ANTHOCYANIDIN 3-O-GLUCOSYLTRANSFERASE 5-LIKE, and ANTHOCYANIDIN SYNTHASE. These enzymes are involved in defense against abiotic and biotic stresses (Landi *et al*., 2015; Zhang & Liu, 2015; Qiu *et al*., 2016; Jin *et al*., 2019).

**Figure 6.**
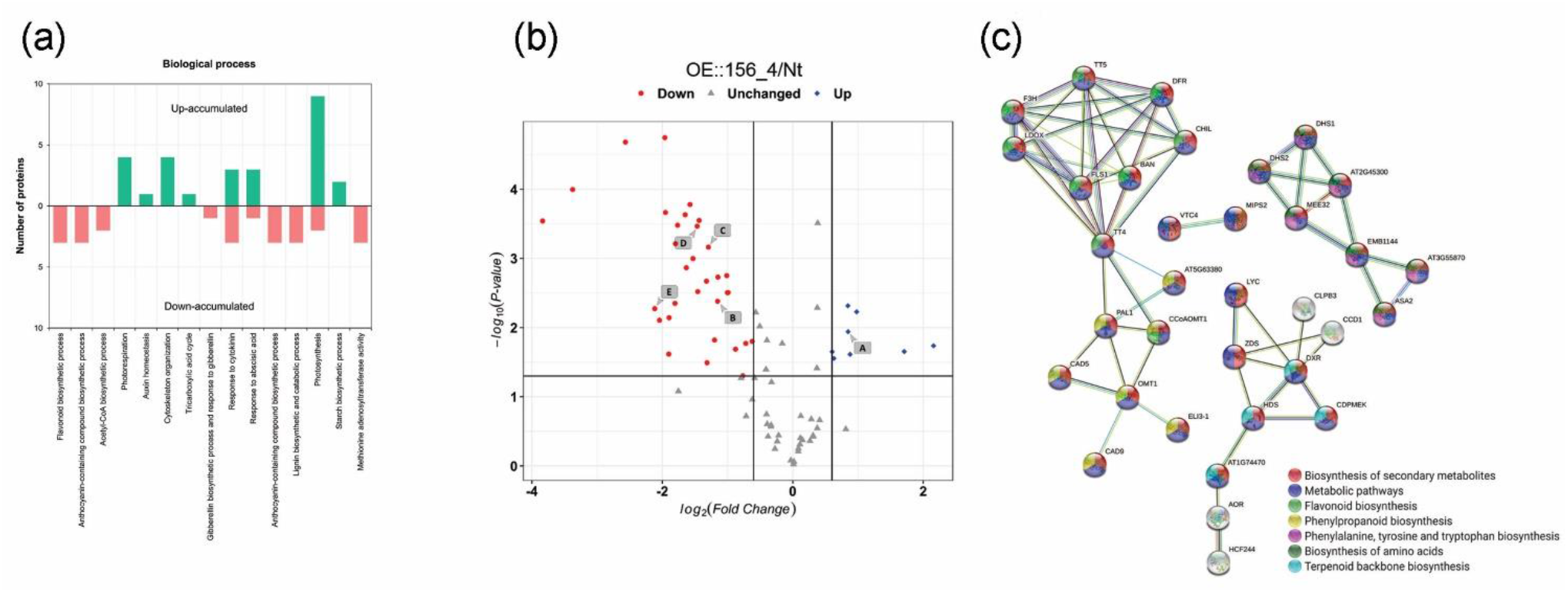
Proteins associated with secondary metabolism are differently accumulated in OE::156 leaves. **(a)** Functional annotation of differentially altered proteins. Biological processes were enriched via Fisher’s exact test with *P* = 0.01 to compare proteins in miR156-overexpressing (OE::156_4) and Non-transformed (Nt) leaves. **(b)** Volcano plot for identifying secondary metabolism-related proteins. Each point represents the difference in expression (fold-change) between OE:: 156_4 and Nt annatto plants plotted against the level of statistical significance. Red dots: down-regulated proteins; gray dots: proteins whose levels have not changed; blue dots: up-regulated proteins; A: ZEAXANTHIN EPOXIDASE; B: FLAVONOL SULFOTRANSFERASE-LIKE; C: PHENYLALANINE AMMONIA-LYASE; D: ANTHOCYANIDIN 3-O-GLUCOSYLTRANSFERASE 5-LIKE; E: ANTHOCYANIDIN SYNTHASE. **(c)** STRING protein-protein interaction diagram. Protein-protein interaction network from the OE::156 and Nt comparison with putative roles in secondary metabolism. The network view summarizes the network of predicted associations for a particular group of proteins based on *Arabidopsis* orthologs identified by STRING using the *B. orellana* amino acid sequence. Only first neighbors are shown in the interactions. All the predicted interactions in the network showed high confidence (> 0.7) according to the combined STRING score. Different colors indicate different KEGG pathways. The straight line shows the interaction between proteins. Yellow line: literature evidence; green line: neighborhood evidence; purple line: experimental evidence; light blue line: database evidence; and black line: co-expression evidence.

Proteins shown in Table S5 were used to determine their mutual functional relationship, which identified four different groups (Fig. 6c): Group I (TT5, DFR, F3H, LDOX, FLS1, BAN, CHIL, and TT4), which are associated with the biosynthesis of amino acids, and with general secondary metabolism; Group II (JAT, PAL1, CCoa, CAD5, OMT1, ELI3-1, and GAD9) linked to the biosynthesis of phenylpropanoid and lignin; Group III (LYC, ZDS, DXR, HDS, CDPMEK, and CCD1) related to carotenoid and ABA biosynthesis, classified also as terpenoid backbone biogenesis; and Group IV (DHS2, DHSI, AT2G, MEE32, EMB1144, AT3G, and ASA2), which contains enzymes implicated in the biosynthesis of specific amino acids. As revealed by the composition of Group III, the main enzymes related to carotenoid biosynthesis were associated together (Fig. 6c). Thus, ABA and carotenoid biosynthesis in leaves was affected by the same factor (i.e., loss of *BoSPL* activity), most likely via the *BoCCD1* or *BoCCD4-4* pathway, thereby reinforcing that the carbon flux was diverted from bixin to ABA production.

**Figure S4.**
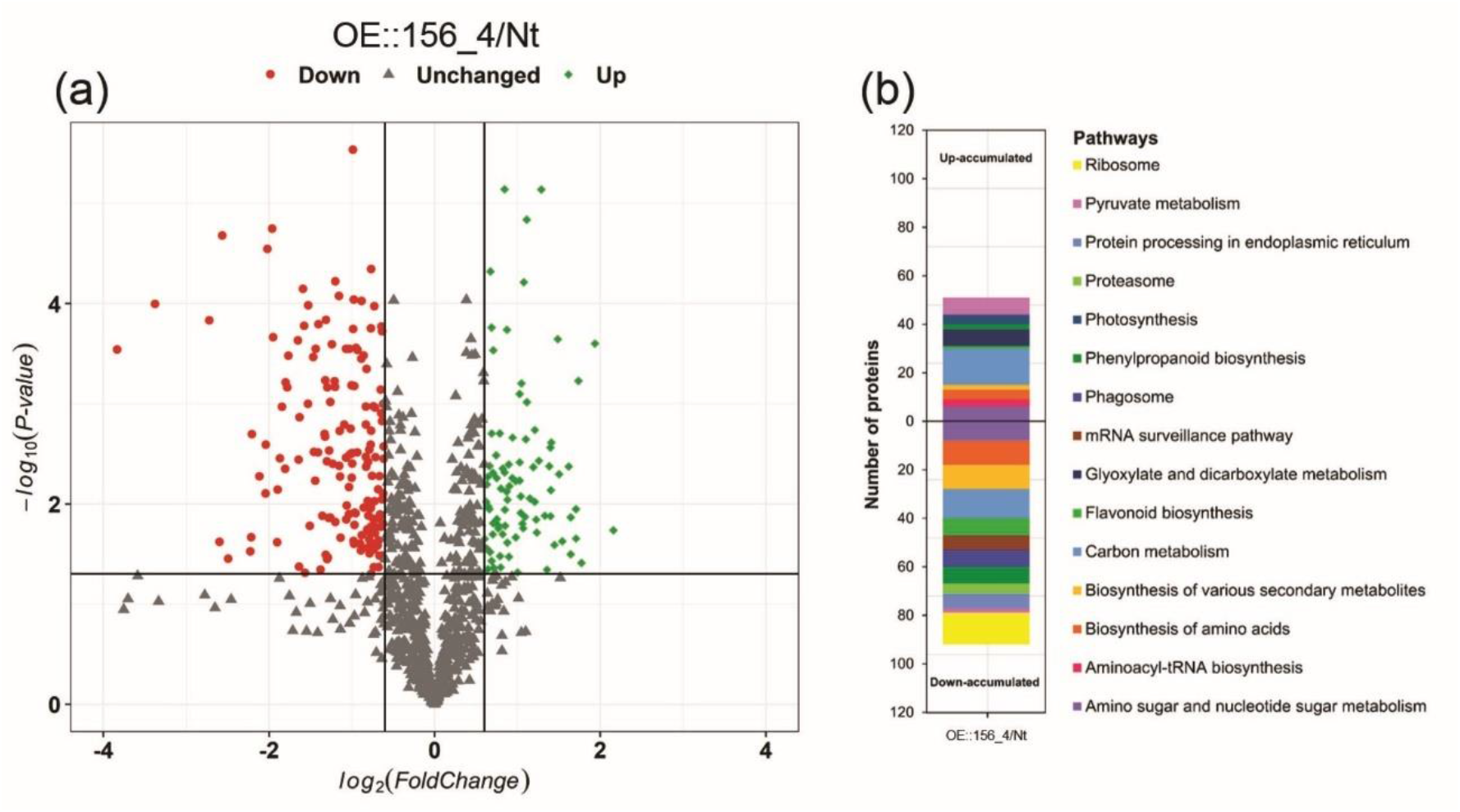
Proteomic analysis of annatto miR156-overexpressing (OE::156_4) and Non-transformed (Nt) leaves. **(a)** Volcano plot of the entire set of proteins quantified in this study. Each point represents the difference in expression (fold-change) between OE::156_4 lines and Nt leaves plotted against the level of statistical significance. Red dots: down-regulated proteins; gray dots: proteins whose level did not change; green dots: up-regulated proteins. **(b)** KEGG pathway enrichment analysis (*P* ≤ 0.01) of differentially accumulated proteins (DAPs) identified from the OE:: 156_4/Nt comparison.

## Discussion

### *BoSPL* activity is required for *B. orellana* vegetative phase change

*B. orellana* plants show few obvious vegetative modifications from juvenile to adult developmental stage, as described by Baliane (1982), which makes it challenging to determine modifications during leaf and shoot development. Determining how phase-specific traits contribute to *B. orellana* fitness is fundamental for better management of this economically important tree. Here, we characterized juvenile annatto trees overexpressing the miR156, and found that modifications during vegetative phase change are not very different from those observed in *Arabidopsis*, an annual herb. miR156-overexpressing *B. orellana* plants exhibited smaller leaf blades shaped as triangles rather than hearts (as commonly displayed by adult *B. orellana* plants; Baliane, 1982), lack of extrafloral nectaries, and thinner bark layer compared to control plants. In poplar *(Populus* sp.), another woody species with subtle vegetative phase changes, the overexpression of miR156 also leads to more evident shoot modifications (Rubinelli *et al*., 2013; Lawrence *et al*., 2020). Collectively, these observations indicate that vegetative phase change is an ancient and fundamental aspect of shoot development, and it might be under natural selection (Leichty & Poethig, 2019).

The higher number of leaves observed in OE::156 annatto plants is a common phenotype observed in *SPL*-deficient plants, such as *Petunia* and tomato (Silva *et al*., 2014; Zhou *et al*.,2021). Smaller leaf size, which indicates modifications in leaf ontogeny, might be a result of differential modulation of genes associated with organ size that are regulated by the miR156/*SPL* module (Zhang *et al*., 2015; Fouracre & Poethig, 2019; Fang *et al*., 2021; Barrera-Rojas *et al*.,2020). Even though OE::156 lines have more leaves, they exhibited similar photosynthetic area as their Nt counterparts, a phenomenon also observed in *Eucalyptus grandis* (Levy *et al*., 2014), *S. tuberosum* (Bhogale *et al*., 2014), and *Arabidopsis* (Xie *et al*.,2017). High levels of miR156, together with the lack of *BoSPLs* and BomiR172 (Fig. 4), suggested that even under greenhouse conditions, OE::156 annatto plants displayed similar overall gene regulation as *in vitro*-raised plants (Faria *et al*., 2022). Importantly, BomiR172 was less abundant in OE::156 leaves, which might correlate with their loss of leaf cell maturation, as observed in rice and *Arabidopsis* (Zhu *et al*., 2009; Jung *et al*., 2011).

### *BoSPL* activity is required for bixin biosynthesis in *B. orellana* leaves

OE::156 leaves exhibited less bixin channels, which likely impacted bixin content (Fig. 2). Bixin gland ontogenesis depends on the gradual coalescence of cells with shared cell walls, which become degraded as seen in laticifer channels (Almeida *et al*., 2021). Such process seems to depend on leaf maturation (Lee and Ding, 2016), which is delayed in OE:: 156 leaves. However, bixin biosynthesis-associated enzymes are also sensitive to changes in the level of miR156.

Transcriptome data revealed differential expression of several genes associated with bixin biosynthesis during *B. orellana* seed development (Moreira *et al*., 2022). Although this study lacks transcriptome data from annatto leaves, by combining gene expression and proteomic analyses, we shed light on how the miR156-targeted *BoSPLs* regulate carotenoid- and bixin-associated pathways at the molecular level (Fig. 7). For instance, the overexpression of miR156 attenuated the transcription of *BoCCD4-4*, which encodes enzyme responsible for lycopene cleavage to bixin aldehyde during bixin biosynthesis. This enzyme is conserved in most eudicots and plays a central role in apocarotenoid production (Varghese *et al*., 2021; Brandi *et al*., 2011). As expected, genes coding for lycopene-associated downstream enzymes were down-regulated in OE::156 plants, except for *BoALDH3I1*(Fig. 5), whose up-regulation could be explained by the *BoALDH3* expression being controlled via the ABA stress response pathway rather than directly by the miR156/*SPL* module (Brocker *et al*., 2013). The *ALDH* gene family has additional functions in plants, as shown in *Arabidopsis*, where ALDHs play a central role in salt tolerance through putrescine and γ-aminobutyrate production (Zarei *et al*., 2016). Loss of *BoSPL* activity in OE::156 leaves led to the up-regulation of enzymes involved in *β*-carotenoid production (Fig. 5). Interestingly, *BoCCD1* up-regulation in OE::156 leaves might be responsible for diverting carbon from bixin to ABA production (Cárdenas-Conejo *et al*., 2015;Cazzonelli & Pogson, 2010). Transcripts of *B. orellana LYCOPENE B-CYCLASES 1* and *2 (Boβ-Lyc1* and *Boβ-Lyc2)* accumulated at higher levels in OE:: 156 leaves compared with Nt counterparts (Fig. 5). MiR156-targeted *MuSPL16* is reported to control secondary metabolism in banana, directly promoting *LYCOPENE B-CYCLASE* activity and thereby increasing β-carotene content in the fruits (Zhu *et al*., 2020). Here, we showed that the loss of *BoSPL* activity seems to increase *LYCOPENE B-CYCLASE* gene expression in leaf tissue (Fig. 5). It is possible that *BoSPLs*, such as *BoSPL6* and/or *BoSPL13*, might directly control annatto *β-Lyc1* and *β-Lyc2* expression to modulate carotenoid and ABA biosynthesis in leaves. This conjecture deserves further investigation.

**Figure 7.**
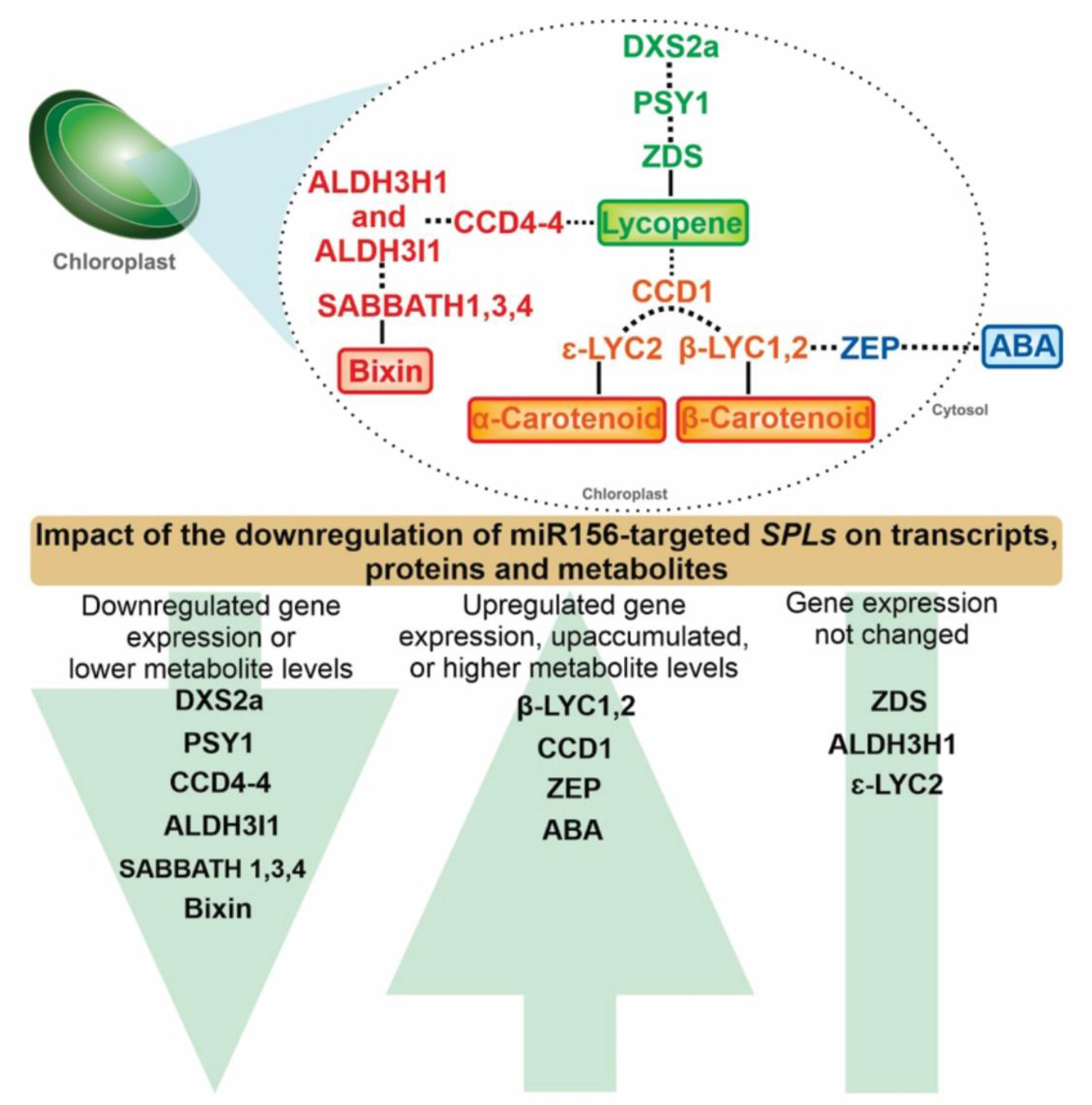
The miR156/*SPL* module controls bixin and ABA biosynthesis. General view of carotenoid-related transcripts, proteins/enzymes, and metabolites changed by the overexpression of miR156 in *B. orellana* plants. Dotted line: indirectly related to the route; continuous line: directly related in the metabolic pathway. Proteins/enzymes names as described in Fig. 5.

Given that bixin and ABA contents were altered in OE::156 leaves (Fig. 2, 5), we investigated whether additional secondary metabolic pathways were affected by the overexpression of miR156. LC-MS analysis revealed that the abundance of several secondary metabolism-associated proteins was reduced in OE::156 leaves (Fig. S4a), with a few exceptions such as SUPEROXIDE DISMUTASE, which responds to high ABA levels (Zhang *et al*., 2008; Salazar-Chavarría *et al*., 2020). Proteins/enzymes involved in both primary and secondary metabolic pathways might be directly or indirectly regulated by the miR156/SPL module, as shown by our proteomic data. Protein/enzyme levels are good indicators of metabolite accumulation because they avoid post-transcriptional regulation, and thus, directly reflect how biochemical pathways respond to stimuli or regulatory routes. In terms of secondary metabolic pathways, our proteomic data revealed that proteins associated with flavonoid, anthocyanin, and lignin biosynthesis were the most affected by the reduction in miR156-targeted *BoSPL* activity (Fig. 6). One of the most interesting enzyme was ZEP, which is crucial for ABA biosynthesis (Marin *et al*., 1996; Hieber *et al*., 2000). However, a recent study suggests a new ABA biosynthesis pathway independent from ZEP, which starts upstream from 9-cis-zeaxanthin (Jia *et al*., 2022). Importantly, the accumulation of ZEP in OE:: 156 leaves suggests that the carbon flux going to bixin production might be rerouted toward ABA biosynthesis during the juvenile phase. There are relatively few examples of juvenile and adult traits, such as secondary metabolites, that might contribute to plant fitness or expected to be selectively advantageous (Leichty and Poethig, 2019; Lawrence *et al*., 2021). Our observation that *B. orellana* adult leaves produce more bixin and less ABA than juvenile ones suggest that may be advantageous to accumulate stress-related metabolites, such as ABA, early in development, when annatto trees are more susceptible to biotic and abiotic stresses. In contrast, increased apocarotenoids as bixin in leaves may be advantageous once adult plants are established and less susceptible to stresses.

The miR156/*SPL* module has been well characterized in numerous species and is highly conserved in angiosperms (Morea *et al*., 2016). However, how different metabolites and their developmental features are controlled by this miRNA regulatory module is still unclear. Here, we illustrate one of these fine-tuned controls of secondary metabolic pathways in *B. orellana*, an economically important species whose molecular characterization is still in its infancy.

## Supporting information

Table S1. Oligonucleotides used in this study.

Table S2. List of SPL code gene and their respective gene name for each plant species analyzed.

Table S3. Total content of chlorophyll a (Chl a) and b (Chl b), carotenoids, and anthocyanin adjusted by fresh weight in non-transformed (Nt) and OE::

Table S4. List of proteins identified in OE::156_4 and Nt plants, and catalogued by Gene Ontology function analysis.

Table S5. List of secondary metabolism-associated proteins extracted from the proteome list.

## Data availability

Data sharing not applicable to this article as no datasets were generated or analyzed during the current study.

## Acknowledgements

The Fundação de Amparo à Pesquisa do Estado de Minas Gerais – FAPEMIG (Grant numbers APQ-02372-17 and APQ-00772-19), the Conselho Nacional de Pesquisa e Desenvolvimento Científico e Tecnológico (CNPq), and Coordenação de Aperfeiçoamento de Pessoal de Nível Superior (CAPES) (Finance Code 0001) are acknowledged for their financial support. Chr. Hansen Ind. Com. Ltda. (Valinhos, SP, Brazil) is acknowledged for kindly donating the seeds of *Bixa orellana* Piave Vermelha, and Dr. A. R. Leichty for kindly making available the miR156 overexpressing vector. K.L.G.M. received a scholarship from CNPq (Grant number 146061/2019-5). D.V.F. received a postdoctoral scholarship from CNPq (Grant number 155615/2018-1). We would like to thank the Universidade Federal de Viçosa (UFV) for providing all relevant infrastructure, to the Núcleo de Análise de Biomoléculas (NUBIOMOL), for ABA and Bixin quantification, to Gilmar Valente from Núcleo de Microscopia e Microanálises (NMM) for microscopy analyzes, to the Genética Molecular de Bactérias Laboratory (DMB – BIOAGRO) for RT-qPCR equipament, and Editage (www.editage.com) for English language editing.

## Author contributions

K.L.G.M., M.B.S.D., D.V.F., and W.C.O. designed the study. K.L.G.M., D.V. F., M.B.S.D., T.F., and E.R. performed the experiments. K.L.G.M., D.V.F., T.R.O., E.R., and W.C.O. analyzed the data. L.A.S.S., T.R.O., D.V.F., W.C.O., and F.T.S.N. provided technical assistance. K.L.G.M., L.A.S.S., D.S.B., D.V.F., M.G.C.C., C.S.C., V.S., E.R., W.C.O., and F.T.S.N. wrote the manuscript.

## Supporting Information

**Table S1. Oligonucleotides used in this study.**

**Table S2. List of *SPL* code gene and their respective gene name for each plant species analyzed**.

**Table S3. Total content of chlorophyll *a* (Chl *a*) and *b* (Chl *b)*, carotenoids, and anthocyanin adjusted by fresh weight in non-transformed (Nt) and OE::156 (OE::156_1 and OE::156_4) leaves of *Bixa orellana* plants.**

**Table S4. List of proteins identified in OE::156_4 and Nt plants, and catalogued by Gene Ontology function analysis.**

**Table S5. List of secondary metabolism-associated proteins extracted from the proteome list.**

